# Exposure of male mice to nicotine leads to metabolic dysfunction in their male and female offspring

**DOI:** 10.1101/2025.10.04.680466

**Authors:** Stephanie Aguiar, Truman Natividad, Daniel Davis, Carlos Diaz-Castillo, Raquel Chamorro-Garcia

**Author notes:** Corresponding author contact information: Raquel Chamorro-Garcia, PhD, University of California, Santa Cruz, Department of Microbiology and Environmental Toxicology, 1156 High Street, Santa Cruz, Ca, 95064. **Competing Interests:** The authors have nothing to disclose.

## Abstract

Paternal exposure to nicotine has been linked to altered metabolic phenotypes in offspring, yet the underlying mechanisms and sex-specific outcomes remain elusive. In this study, we investigated the effects of paternal nicotine exposure on glucose metabolism, and adipose tissue and liver transcriptomic profiles in male and female offspring. Despite no differences in body weight during early postnatal life, female offspring displayed lower fasting glucose and reduced glucose levels during glucose tolerance tests, without alterations in insulin sensitivity. These effects were accompanied by decreased circulating insulin and upregulation of genes involved in the insulin signaling pathway in gonadal white adipose tissue (gWAT), consistent with enhanced glucose uptake capacity. In contrast, male offspring showed no overt changes in glucose tolerance or insulin sensitivity, despite reduced plasma insulin and glucagon levels. In male offspring, liver transcriptomic analyses revealed downregulation of glucagon signaling, insulin resistance, and PPARα pathways, suggesting impaired fasting adaptation and reduced hepatic catabolic capacity. Collectively, these findings demonstrate that paternal nicotine exposure induces sex-dependent metabolic reprogramming, with pronounced effects on glucose homeostasis in females and transcriptional signatures of reduced fasting resilience in males. These results identify the liver and adipose tissue as key targets of paternal nicotine exposure in the offspring and underscore the need for longitudinal studies to determine whether these alterations predispose offspring to metabolic disease across the lifespan.

## INTRODUCTION

The rise in metabolic diseases like obesity, type 2 diabetes (T2D), and non-alcoholic fatty liver disease (NAFLD) has prompted research on environmental factors contributing to their development (Merrill et al. 2020). While lifestyle factors such as diet and physical activity remain crucial to understand the onset of metabolic diseases, exposure to environmental toxicants has emerged as a significant, albeit often overlooked, risk factor (Carbone et al. 2019). Over the past two decades, a growing body of evidence derived from rodent studies has demonstrated that exposure to environmental toxicants can induce metabolic disease, not only in the individuals directly exposed to them but also in subsequent generations that remain unexposed (Chamorro-García et al. 2013; Chamorro-Garcia et al. 2017; Chamorro-García et al. 2021; Diaz-Castillo et al. 2025; King et al. 2019b, 2019a; Nilsson et al. 2023). Among these environmental exposures, smoking has been identified as a factor that increases susceptibility to metabolic diseases such as obesity, type 2 diabetes, and cardiovascular disease (Akbartabartoori et al. 2006).

Smoking tobacco products is one of the leading preventable causes of adverse health effects, including respiratory diseases, an increased risk of developing type 2 diabetes, and high blood pressure in adults in the United States (Cornelius et al. 2023). Although cigarette smoking has decreased in recent decades (The Health Consequences of Smoking—50 Years of Progress), the advent of electronic cigarettes (e-cigarettes) has led a new generation of young adults to use tobacco products (Erhabor et al. 2023). This approach is deemed safer than the traditional cigarette smoking practice due to the absence of combustion byproducts. However, nicotine, the primary psychoactive component of tobacco, has been recognized as an endocrine-disrupting chemical that can affect thyroid, adrenal, and reproductive axes as well as other risk factors for metabolic disease, such as elevated blood pressure and free fatty acids in plasma, and a decreased mobilization of blood glucose (Lee et al. 2025; Tweed et al. 2012). Despite there is a tendency of higher rates of male smokers than female smokers, with 36.7% and 7.8% smokers, respectively (Siddiqi et al. 2020), most of the efforts to understand the multigenerational effect of tobacco use have been placed in studying maternal exposure to tobacco products during pregnancy.

Investigations into the paternal contributions to the health of subsequent generations have revealed that environmental exposures can induce metabolic alterations in progeny in rodents. Specifically, paternal environmental exposures to suboptimal diets, alcohol, and cocaine have resulted in neurological, behavioral, and physiological alterations in the next generation (Carone et al. 2010; Finegersh and Homanics 2014; Yaw et al. 2022). In humans, paternal nicotine exposure can lead to cognitive deficits and behavioral alterations in the next generation (Maurer et al. 2022). In rodents, paternal nicotine exposure has been known to lead to transcriptomic alterations in their male offspring, including alterations in the expression of genes involved in the metabolism of hepatic lipids, fatty acids, and xenobiotics in a mouse model (Vallaster et al. 2017).

In this study, we investigated the metabolic effects of paternal exposure to nicotine on female and male offspring. Male mice were exposed to nicotine through drinking water and mated to untreated females. We analyzed metabolic endpoints, including body weight, glucose homeostasis, plasma metabolites involved in metabolic regulation, and transcriptomic data from liver and gonadal white adipose tissue (gWAT). In females, we observed significant alterations in glucose homeostasis and liver and gWAT function. In contrast, only significant alterations in liver function were observed in males.

## METHODS

### Chemicals and Reagents

(-)-Nicotine (#N3876-25mL), D-(+)-glucose (#G8270), and human recombinant insulin (dry powder, #91077C) were purchased from Sigma-Aldrich. Dimethyl sulfoxide (DMSO) was purchased from Fisher Scientific, LLC. Nicotine was stored out of light and in a desiccator. Glucose and insulin stocks for glucose and insulin tolerance tests were prepared fresh the day of metabolic testing.

### Animal Maintenance and Exposure

Animals were purchased at Jackson Laboratory (Sacramento, CA). Animals were housed in micro-isolator cages in a temperature-controlled room (21-22°C) with a 12 h light/dark cycle and provided food (Envigo; Teklad Global Soy Protein-Free Extruded Rodent Diet, irradiated #2920X) and water *ad libitum* unless otherwise indicated. Animals were treated humanely and with regard for alleviation of suffering. All procedures conducted in this study were approved by the Institutional Animal Care and Use Committee of the University of California, Santa Cruz Protocol # Chamr1908. All tissue harvesting was performed with the dissector blinded to which groups the animals belonged. At the moment of euthanasia, each mouse was assigned a code, known only to the lab member not involved in dissections.

Three-week-old C57BL/6J male mice (n=20) were purchased from Jackson Laboratory (Sacramento, CA), and randomly assigned to two treatment groups receiving 200 µg/mL nicotine diluted in dimethyl sulfoxide (DMSO) (n=10) or 0.1% DMSO (vehicle control, n=10) in drinking water (Matta et al. 2007). One mouse from the control group had to be euthanized due to malocclusion resulting in only 9 animals in that group. Every 3-4 days, we measured and discarded the remaining water in the bottles, and water bottles were refilled with freshly prepared treated water. The treatment continued for six weeks, to encompass the entirety of spermatogenesis process to ensure that sperm at all stages were being exposed to oral consumption of nicotine (Oakberg 1956). After the sixth week of treatment, each male mice was mated with a single age-matched unexposed C57BL/6J female mice purchased from Jackson Laboratory.

Pregnancies were confirmed by detection of gestational plugs and body weight gain one week after the detection of the plug. F0 males were euthanized upon observation of gestational plug. Litter size and sex ratio were recorded after birth (Supplemetnal Table 1). Since litter size can affect growth trajectories of the pups, we only considered litters that had between 6-8 pups (average litter size in our cohort 6.5-7 pups), and litters with less than 2 members of each sex were excluded. We considered both male and female offspring separately in our analysis.

Ten F1 animals per paternal exposure group and sex were weaned from dams at 3 weeks old and placed on a synthetic diet (Envigo, 93G, TD.140148) for five weeks. Diet was supplemented with fresh pellets every week. Weekly body weight measurements were recorded between weeks 2-8. At 8 weeks old, animals were euthanized via isoflurane overdose followed by cardiac exsanguination. Blood was drawn directly from the heart using EDTA-treated syringe and needles and placed in a clean tube containing protease inhibitors (Protease Inhibitor Cocktail, EDTA-free, Sigma-Aldrich #S8830). Blood was centrifuged for 10 min at 5,000 rpm at 4ºC. Plasma was transferred to a clean tube, snap-frozen in liquid nitrogen and preserved at - 80ºC. Samples were shipped to Eve Technologies Corporation (Calgary, AB) for analysis of a panel of plasma metabolites (Mouse/Rat Metabolic Hormone Discovery Assay® 11-Plex, MRDMET). Liver and gonadal white adipose tissue (gWAT) samples were collected and weighed from animals. Both tissues were snap frozen and stored at -80ºC for RNA sequencing analyses.

### Cotinine levels in plasma

Male C56BL/6J animals were exposed to 200 µg/mL of nicotine or 0.1% DMSO via drinking water for six weeks (8 animals per group). A subset of 4 animals was euthanized via isoflurane overdose and cardiac exsanguination after 3 and 6 weeks of exposure. Blood was processed to obtain plasma as described in the “Animal maintenance and exposure” section. Cotinine levels in plasma were measured via ELISA (Mouse/Rat Cotinine ELISA Kit; Fisher Scientific Company LLC).

### Glucose and insulin tolerance tests

At six and seven weeks of age, five animals per group were subjected to glucose and insulin tolerance tests (GTT and ITT), respectively. Glucose and insulin stocks were prepared fresh the day of the assay in 0.9% saline. After 4 hours of fasting, animals were administered 2 g of glucose/kg body weight (b.w.) or 0.75 IU of insulin/kg b.w. via intraperitoneal injection. Glucose levels were measured with Contour® blood glucose meter and Contour® blood glucose strips every 30 minutes for 120 minutes after injection of glucose or insulin. Upon GTT and ITT completion, mice were returned to their corresponding cages with *ad libitum* access to food.

### RNA Isolation and sequencing

RNA from F1 liver was isolated using Direct-zol RNA MiniPrep (Zymo Research #R2053). Tissues were homogenized with VWR Premium Micro-Homogenizer (#10032-328). RNA from 5 randomly selected non-sibling mice from each group were submitted to the University of California Davis DNA Technologies & Expression Analysis Core Laboratory for 3’ Tag-RNA-sequencing using an Illumina HiSeq 4000 instrument. Sequencing read processing and determination of significant transcriptome variation were performed using Galaxy Project platform (Galaxy version 23.0) (Afgan et al. 2022). FastQ files were processed using FastQC to determine the quality of the sequencing data obtained (Galaxy version 0.73). Cutadapt (Galaxy version 4.9+galaxy1) was used to trim low quality bases. Read Indexing and alignment to the mouse genome (mm39) was done using STAR (Galaxy version 2.7.10b+galaxy3). FeatureCounts (Galaxy version 2.03+galaxy2) function was used to assign uniquely mapped RNA-seq reads to GRCm39 mouse reference genome count reads. DESeq2 (Galaxy version 2.11.40.8+galaxy0) function was used to determine differentially expressed genes between the nicotine and control groups. We deemed expression levels to be statistically significant if p-value was lower than 0.05.

Kyoto Encyclopedia of Genes and Genomes (KEGG) pathway enrichment analysis was performed using the DAVID bioinformatics resource (Huang et al. 2009b, 2009a). The KEGG pathway database is a collection of pathway maps that illustrate the current understanding of molecular interaction, reaction, and network relationships of any given list of genes, proteins or chemical compounds (Kanehisa et al. 2016). SRPlots was used to visualize differential gene expression patterns between treatment groups and controls for selected genes of interest (Tang et al. 2023).

### Statistical analyses

Statistical analyses for metabolic endpoints (body weight, plasma metabolites, and glucose and insulin tolerance tests were performed using GraphPad Prism 10.0 (GraphPad Software, Inc.). Statistical tests and specific comparisons are indicated in each figure and their respective figure legend.

## RESULTS

### F0 Males exposed to nicotine had significantly decreased body weights

Male mice were exposed to 200 µg/mL nicotine in their drinking water for six weeks to ensure exposure was present throughout most of the spermatogenesis cycle (Oakberg 1956). The control group was exposed to 0.1% DMSO, as DMSO is the solvent used for nicotine in this study. The nicotine dosage resulted in blood cotinine levels ranging from 500 to 1000 ng/mL (Supplemental Figure 1A), which are comparable to those found in average smokers (Keskitalo et al. 2009). Male mice exposed to nicotine exhibited a trend towards decreased weekly body weights when compared to vehicle control animals, although the difference was not statistically significant (Supplemental Figure 1B).

### Paternal exposure to nicotine leads to altered glucose metabolism in female offspring

Female offspring of males exposed to nicotine during a six-week period did not exhibit any substantial alterations in body weight (BW) within the first eight weeks (Figure 1A). However, glucose tolerance test (GTT) conducted on the sixth week indicated that females from the nicotine group exhibited lower fasting glucose levels immediately before starting the assay (Timepoint 0) and 30 minutes after glucose injection (Figure 1B). Notably, we did not observed changes during ITT between groups (Figure 1C). Additionally, we analyzed plasma levels of twelve metabolites involved in the regulation of metabolic processes: active amylin, c-peptide 2, total gastric inhibitory polypeptide (GIP), active glucagon-like peptide-1 (GLP-1), active ghrelin, glucagon, insulin, leptin, PP, PYY, resistin, and secretin. We observed a significant reduction in insulin levels in females from the nicotine group when compared to control females. Additionally, we found a slight increase in ghrelin levels, although this latter finding did not reach statistical significance (p=0.052) (Figure 1D-E).

**Figure 1.**
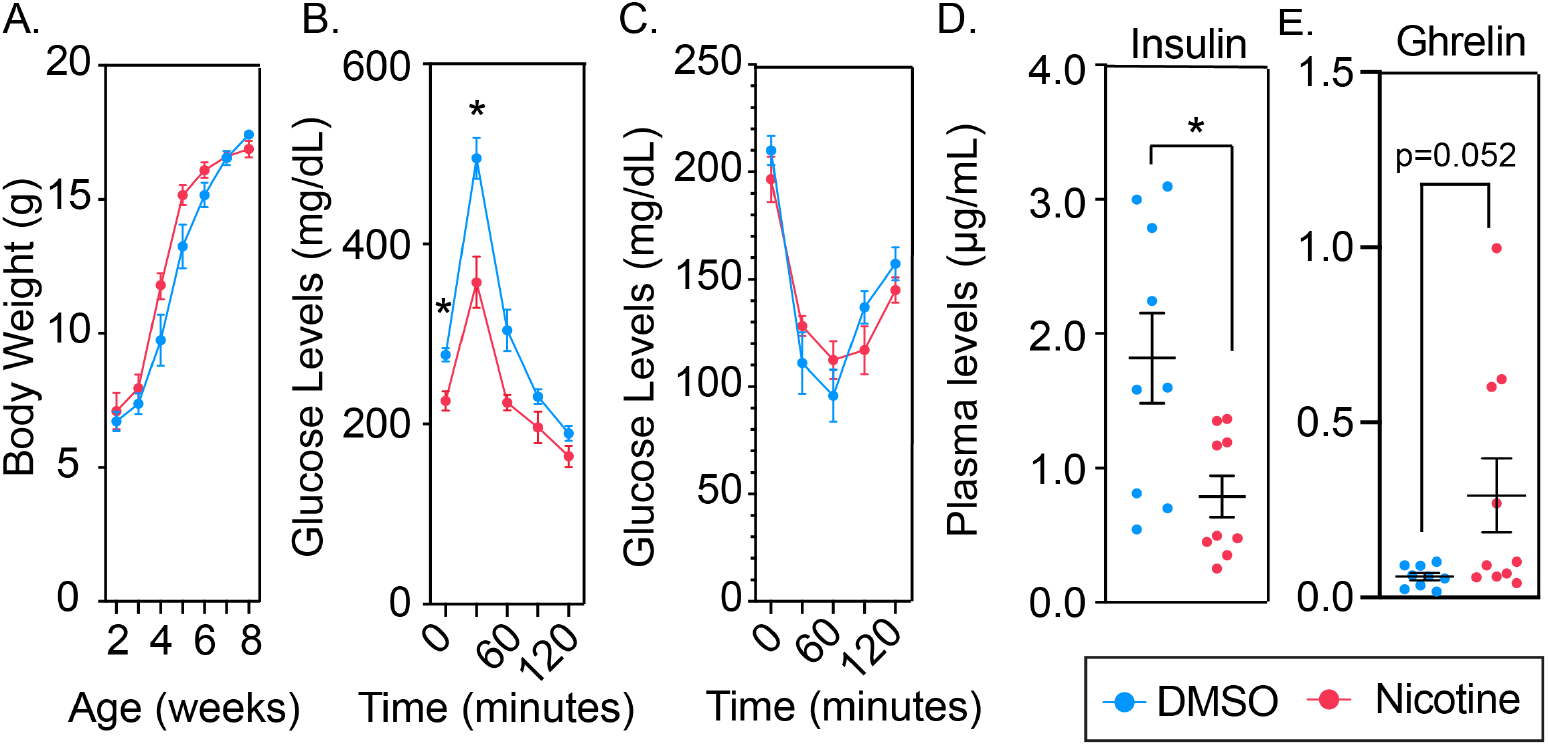
Metabolic assessment of female offspring after paternal exposure to nicotine. **A)** Body weights at 2-8 weeks of age. **B-C)** Glucose and insulin tolerance tests performed at 6 and 7 weeks of age, respectively, after 4 hours of fasting (n=5). **D-E)** Circulating insulin and ghrelin levels at 8 weeks of age after overnight fasting (n=9-10). Statistics: Panels A-C, Two-Way ANOVA followed by Šídák’s post-test; panels D-E, t-test. *p-value <0.05.

To determine significant differences in gene expression caused by paternal exposure to nicotine and the functionality of such variation, RNA was isolated from liver and gWAT and performed 3’Tag RNA sequencing. Rather than focusing on individual candidate genes that might be contributing to phenotype described, we decided to focus on the functionality of genes with the most significant difference in expression between nicotine and control samples. To perform functional enrichments of genes differentially expressed, we selected all genes with p-value <0.05 regardless of their log2 fold change and utilize KEGG pathways to determine whether genes that were deemed differentially expressed were statistically overrepresented in any KEGG pathway compared to what would be expected by chance (Table 3). Transcriptomic analyses of gWAT revealed substantial alterations in multiple KEGG pathways. Notably, genes associated to KEGG Insulin Signaling Pathway (mmu04910), such as insulin receptor (Insr) was significantly overexpressed in the nicotine group when compared to controls (Figure 2A-B). Although we did not detect altered plasma glucagon levels, the KEGG Glucagon Signaling Pathway appeared to have a large number of genes overexpressed in the nicotine group compared to the control group in gWAT (Figure 2A, C). Other metabolic pathways that appear significantly altered in female gWAT are thermogenesis (mmu04714) and fatty acid metabolism (mmu01212) (Figure 2A, D, E).

**Figure 2.**
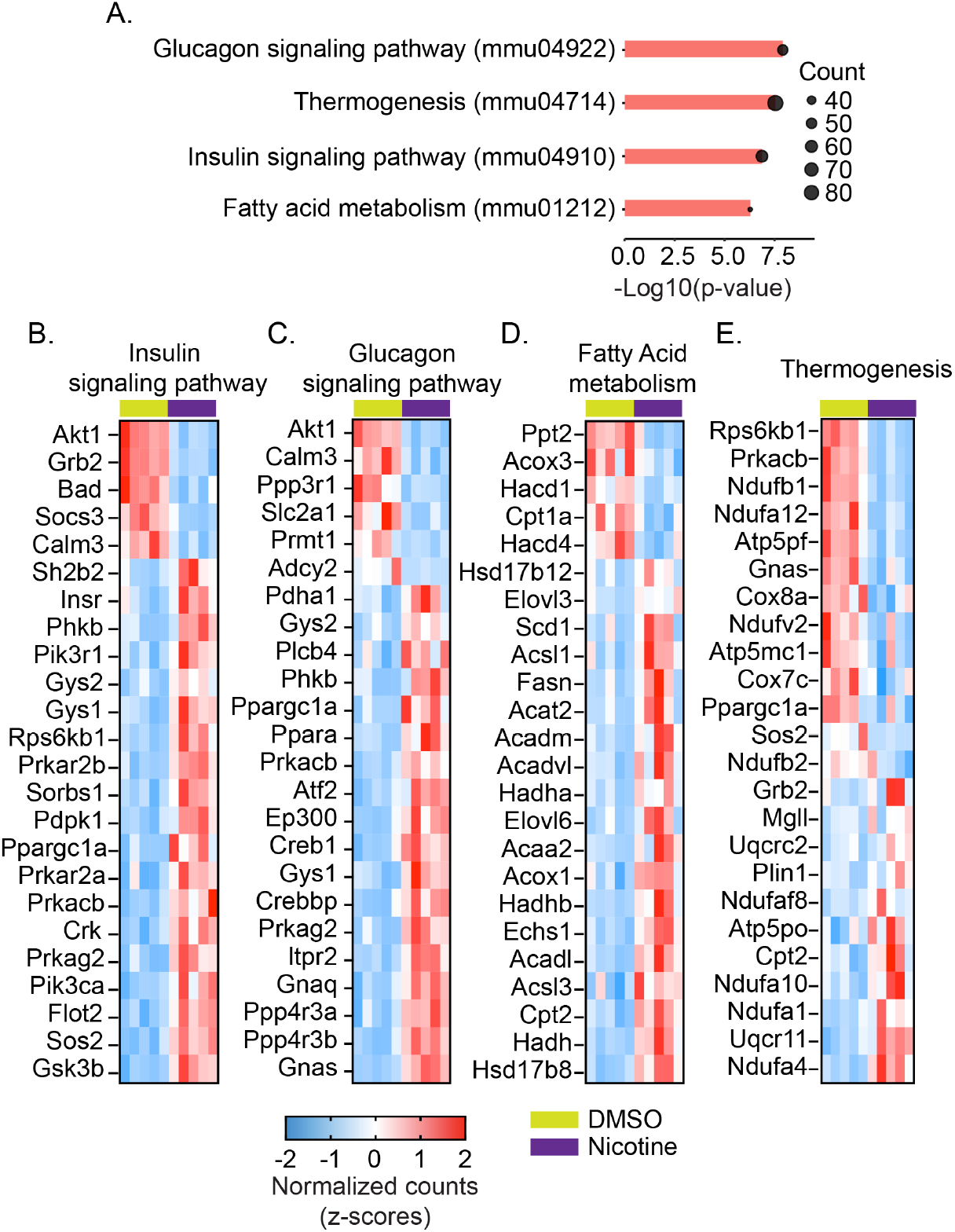
Functional analyses of gWAT in female offspring after paternal exposure to nicotine. **A)** Enriched KEGG pathways associated with metabolic processes revealed by DAVID bioinformatics functional analyses. Log10(p-value) is represented in the x-axis. Count shows the number of differentially expressed genes in each KEGG pathway in nicotine group compared to the control group. **B-E)** Heatmaps representing the top 24 genes with lowest p-value determined with DESeq2 tool in Galaxy Project (z-scores from normalized counts). P-value <0.05 was deemed significant.

We also analyzed transcriptomic alterations in the liver. One of the key roles the liver has is the biosynthesis of glucose from non-carbohydrate carbon substrates including triglycerides, certain amino acids and lactate, which is known as gluconeogenesis. Low levels of glucose during fasting suggest a potential dysfunction in liver gluconeogenesis; however, we did not find enrichment of the KEGG Gluconeogenesis/glycogenesis pathway (mmu04922). In contrast, we found enrichment of KEGG pathways associated with other metabolic processes in the top 10 with lowest p-value, including Metabolic pathways (mmu01100), Cholesterol metabolism (mmu04979) and PPAR signaling pathway (mmu03320).

### Paternal exposure to nicotine leads to alterations in hepatic function in male offspring

In male offspring, we observed no significant alterations in body weight or glucose homeostasis, determined using GTT and ITT, between the nicotine and the DMSO groups (Figure 3A-C). We further analyzed the same plasma metabolic panel as for female offspring. Notably, plasma glucagon and insulin levels were significantly lower in the nicotine group compared to the control group (Figure 3D-E).

**Figure 3.**
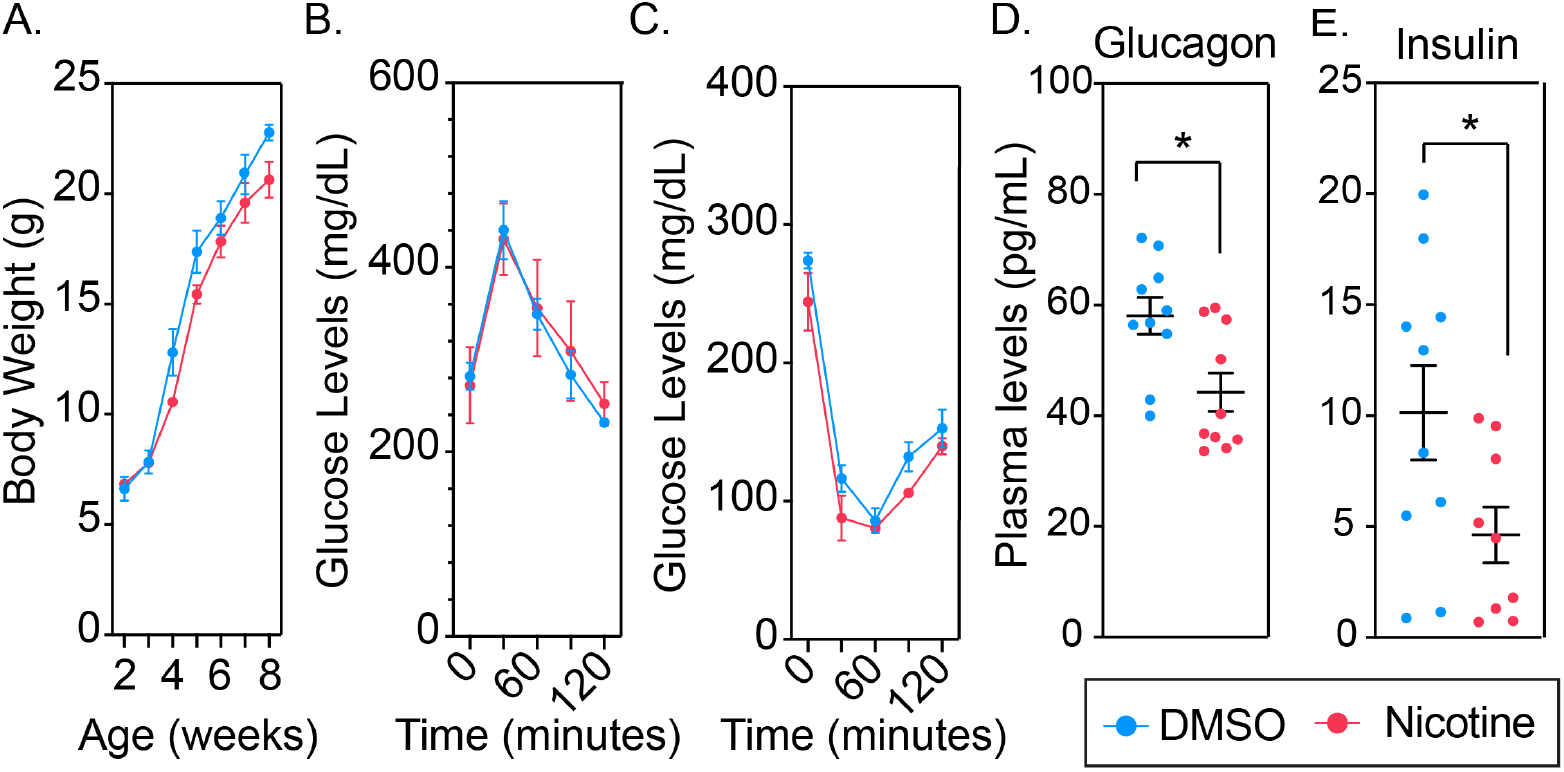
Metabolic assessment of male offspring after paternal exposure to nicotine. **A)** Body weights at 2-8 weeks of age. **B-C)** Glucose and insulin tolerance tests performed at 6 and 7 weeks of age, respectively, after 4 hours of fasting (n=5). **D-E)** Circulating glucagon and insulin levels at 8 weeks of age after overnight fasting (n=9-10). Statistics: Panels A-C, Two-Way ANOVA followed by Šídák’s post-test; panels D-E, t-test. *p-value <0.05.

Transcriptomic analyses of liver tissue showed a significant representation of genes differentially expressed in the KEGG Glucagon Signaling Pathway (mmu04922) and the Insulin Resistance Pathway (mmu04931) in the nicotine group compared to the control group (Figure 4A-C). Underexpression of genes involved in the glucagon and insulin signaling pathways after prolonged fasting (∼16 hours) suggests an impaired catabolic response, potentially compromising the organism’s ability to maintain glucose and energy balance during nutrient scarcity. Additionally, we found that many genes from the KEGG PPAR pathway, including PPAR alpha (*Ppara*), were underexpressed in the liver of the nicotine group compared to the control group. Transcriptomic analyses of gWAT did not reveal significant alterations in gene expression associated with metabolic pathways in the nicotine group compared to the control group.

**Figure 4.**
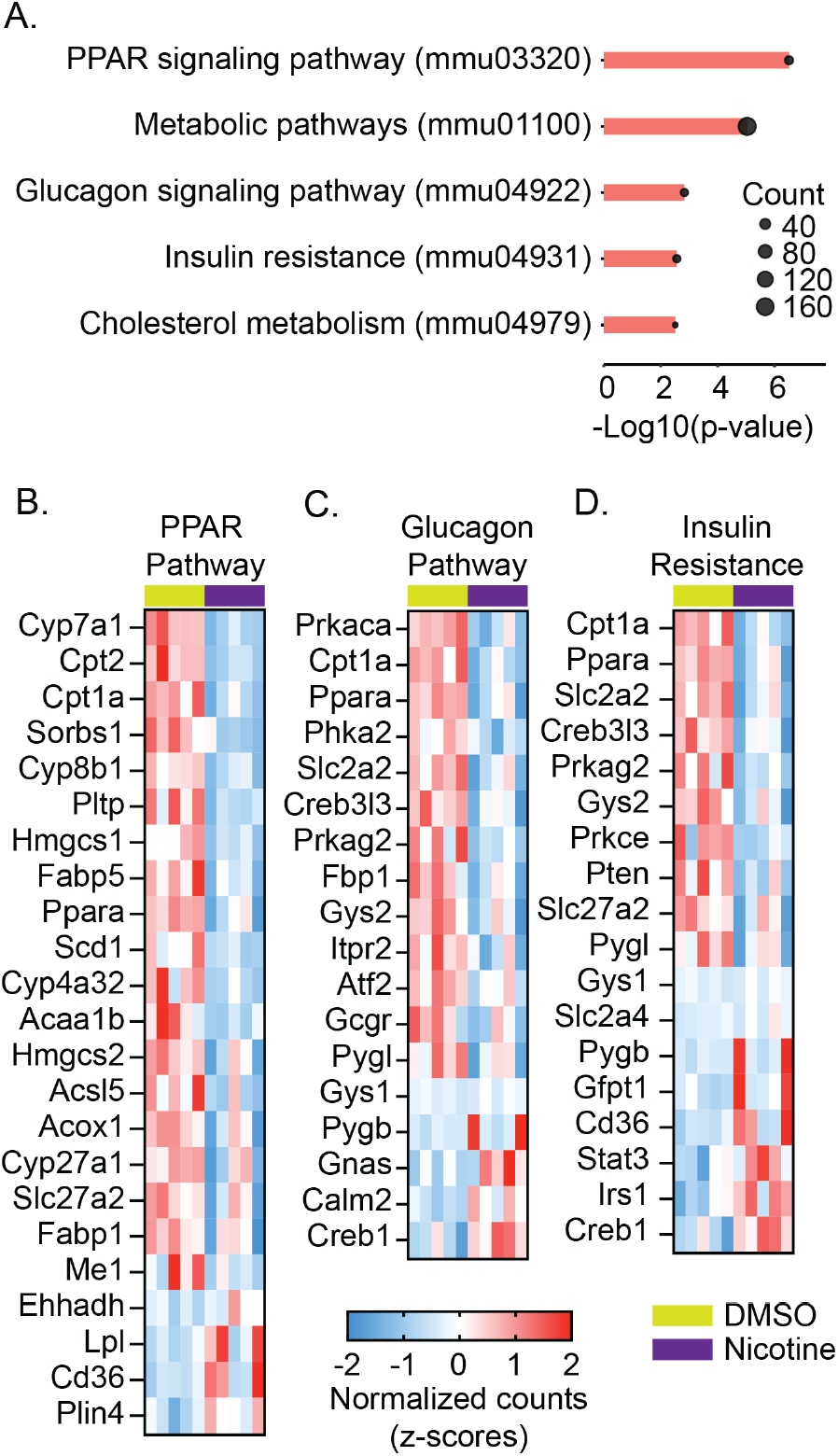
Functional analyses of liver in male offspring after paternal exposure to nicotine. **A)** Enriched KEGG pathways associated with metabolic processes revealed by DAVID bioinformatics functional analyses. Log10(p-value) is represented in the x-axis. Count shows the number of differentially expressed genes in each KEGG pathway in nicotine group compared to the control group. **B-E)** Heatmaps representing differentially expressed genes from the corresponding pathway (z-scores from normalized counts). P-value determined with DESeq2 tool in Galaxy Project. P-value <0.05 was deemed significant.

## DISCUSSION

It is widely recognized that exposure to nicotine can elevate the risk of various diseases, including cardiovascular disease and cognitive impairment, in individuals who consume it (Dorotheo et al. 2024; Wang et al. 2022). Notably, similar effects have been observed in the offspring of pregnant women exposed to nicotine (Bruin et al. 2010). Previous research has demonstrated that paternal exposure to nicotine leads to enhanced drug metabolism, disruption of cognitive function and metabolic disease in their offspring (Li et al. 2016; McCarthy and Bhide 2021; Vallaster et al. 2017). Although there is some literature showing the association between paternal nicotine exposure and offspring metabolic disease (Liu et al. 2022; Vallaster et al. 2017), the metabolic phenotype remains largely uncharacterized. In this study, we investigate the metabolic effects of paternal exposure to nicotine on male and female offspring.

Our findings demonstrate that paternal exposure to nicotine induces sex-specific alterations in glucose metabolism and metabolic signaling in the offspring, despite no major differences in body weight. In female offspring, we observed lower levels of fasting glucose and fast mobilization of glucose during GTT. Interestingly, these effects were not accompanied by changes in insulin sensitivity as measured by ITT, suggesting that the glycemic profile is not attributable to systemic changes in insulin responsiveness, but may reflect tissue-specific alterations in glucose uptake or insulin production by the pancreas. The observed lower fasting glucose levels prior to the GTT but not the ITT indicate a certain degree of variability in the fasting response, potentially attributed to a metabolic adaptation to intermittent fasting. Despite this variation, metabolite analyses conducted on plasma isolated post-mortem after overnight fasting revealed a substantial reduction in insulin levels and a near-significant increase in ghrelin levels in female participants from the nicotine group compared to the control group. These findings align with hormonal responses observed in pathological conditions such as type 1 diabetes and anorexia nervosa (Schalla and Stengel 2018). The transcriptomic analysis of female gWAT provides additional insights into the molecular mechanisms underlying these observations. Specifically, we observed increased expression of genes in the KEGG Insulin Signaling Pathway, which collectively suggest an enhanced insulin responsiveness and glucose uptake capacity in gWAT. Notably, we observed an absence of enrichment of the gluconeogenesis/glycogenesis pathway in the liver of female offspring. Low fasting glucose levels are indicative of impaired hepatic glucose production, yet our transcriptomic results did not support this conclusion. Instead, we observed enrichment of pathways associated with cholesterol metabolism, PPAR signaling, and global metabolic processes, suggesting that paternal nicotine exposure reprograms liver function toward altered lipid handling rather than directly impairing gluconeogenesis. The absence of data regarding glucose levels post-overnight fasting right before euthanasia hinders our ability to discern whether fasting duration exacerbates glucose levels in the nicotine group relative to the control group or if an alternative mechanism is available to promote glucose release into the bloodstream.

In contrast to the female phenotype, male offspring did not show significant differences in glucose tolerance, insulin sensitivity, or fasting glucose levels, despite similar reductions in plasma insulin levels. The lack of alterations in the male mice ability to metabolize glucose may indicate that the organism is more effective in maintaining normoglycemia, or that the primary effects of paternal nicotine exposure in males manifest in other metabolic targets not assessed here, such as lipid handling. The most striking finding in males was the significant reduction in circulating glucagon levels. Glucagon is secreted by the pancreas in response to low glucose levels to stimulate liver gluconeogenesis and inhibit glycolysis, resulting in glucose release into the bloodstream. Low levels of glucagon would suggest a reduction of glucose release from the liver during fasting; however, we did not observe significant changes of fasting glucose. The transcriptomic data from male liver tissue showed an overall underexpression of genes in the KEGG Glucagon Signaling and Insulin Resistance pathways, suggesting a blunted ability to mobilize energy during fasting conditions. Furthermore, the PPAR signaling pathway, including the gene *Ppara*, was downregulated in the liver of male animals from the nicotine group. Reduced PPARα activity may compromise the metabolic shift required during nutrient scarcity, potentially predisposing these animals to hepatic steatosis or impaired adaptation to prolonged fasting (Wang et al. 2020). Such transcriptional changes are consistent with a metabolic phenotype that could increase susceptibility to metabolic disease later in life, including insulin resistance and type 2 diabetes.

Taken together, these findings underscore that paternal nicotine exposure leads to alterations in metabolic networks in the offspring, with functional consequences that are more pronounced in females for glucose handling but may manifest in males as latent susceptibility to metabolic dysfunction. The lack of agreement between transcriptional signatures and metabolic outcomes in some cases (e.g., reduced glucagon but normal fasting glucose in males during short fasting) highlights the need for dynamic metabolic flux studies to clarify the physiological significance of these molecular changes. Future work should focus on assessing the long-term metabolic health of these offspring, including susceptibility to diet-induced obesity, insulin resistance, and hepatic steatosis, as well as potential epigenetic mechanisms of intergenerational inheritance.

In conclusion, our study provides evidence that paternal nicotine exposure reprograms glucose metabolism and liver and adipose tissue function in a sex-dependent manner. Female offspring exhibit enhanced glucose clearance, likely mediated by increased gWAT insulin signaling activity, whereas male offspring show transcriptional signatures suggestive of impaired fasting adaptation and reduced hepatic catabolic signaling. These results highlight the liver and adipose tissue as key targets of paternal nicotine exposure and suggest that both sexes undergo a differentially expressed metabolic reprogramming, with potential implications for disease risk across the lifespan.

## Acknowledgments

This work is supported by the UC Santa Cruz Start-up Funds to Raquel Chamorro-Garcia.

## Data Availability

Transcriptomic data will be available upon acceptance of this manuscript

---

**Supplemental Figure 1.**
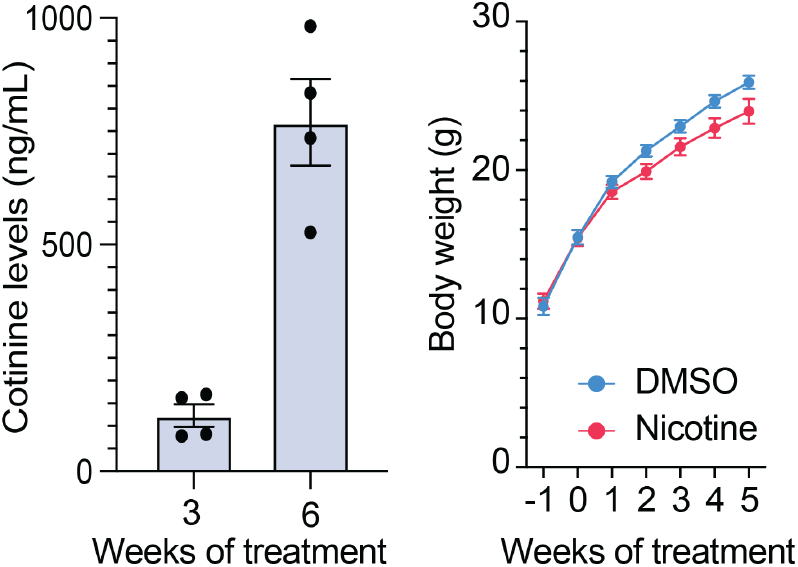
A) Cotinine levels in plasma after 3 and 6 weeks of exposure to 200 µg/mL of nicotine in male mice (n=4). B) Weekly body weight of male C57BL/6J mice exposed to 200 µg/mL of nicotine or 0.1% DMSO in drinking water (n=10). No significant changes (p-value <0.05) were observed between animals exposed to nicotine and DMSO controls. Statistics: 2-way ANOVA followed by Šídák’s post-test.

